# Response splicing QTLs in primary human chondrocytes identifies putative osteoarthritis risk genes

**DOI:** 10.1101/2024.11.11.622754

**Authors:** Seyoun Byun, Philip Coryell, Nicole Kramer, Susan D’Costa, Eliza Thulson, Jacqueline Shine, Sylvie Parkus, Susan Chubinskaya, Richard F Loeser, Brian O Diekman, Douglas H Phanstiel

## Abstract

Osteoarthritis affects millions worldwide, yet effective treatments remain elusive due to poorly understood molecular mechanisms. While genome-wide association studies (GWAS) have identified over 100 OA-associated loci, identifying the genes impacted at each locus remains challenging. Several studies have mapped expression quantitative trait loci (eQTL) in chondrocytes and colocalized them with OA GWAS variants to identify putative OA risk genes; however, the degree to which genetic variants influence OA risk via alternative splicing has not been explored. We investigated the role of alternative splicing in OA pathogenesis using RNA-seq data from 101 human chondrocyte samples treated with PBS (control) or fibronectin fragment (FN-f), an OA trigger. We identified 590 differentially spliced genes between conditions, with FN-f inducing splicing events similar to those in primary OA tissue. We used CRISPR/Cas9 to mimic an SNRNP70 splicing event observed in OA and FN-f-treated chondrocytes and found that it induced an OA-like expression pattern. Integration with genotyping data revealed 7,188 splicing quantitative trait loci (sQTL) affecting 3,056 genes. While many sQTLs were shared, we identified 738 and 343 condition-specific sQTLs for control and FN-f, respectively. We identified 15 RNA binding proteins whose binding sites were enriched at sQTL splice junctions and found that expression of those RNA binding proteins correlated with exon inclusion. Colocalization with OA GWAS identified 6 putative risk genes, including a novel candidate, PBRM1. Our study highlights the significant impact of alternative splicing in OA and provides potential therapeutic targets for future research.

## Introduction

Osteoarthritis (OA) is a chronic, debilitating joint disease affecting over 500 million people world-wide^1^. Despite its prevalence, effective treatments to prevent disease progression remain elusive, due in part to our limited understanding of the underlying molecular mechanisms^2^. Genetic factors play a substantial role in OA risk, with an estimated heritability of over 50%^3^. Genome-wide association studies (GWAS) have identified over 100 loci associated with OA risk^4^, yet understanding their functional impact remains challenging as most associated single nucleotide polymorphisms (SNPs) reside in non-coding genomic regions^5,6^.

One powerful method to determine the mechanisms through which disease-associated variants act is to identify quantitative trait loci (QTL) for a feature of interest (e.g. gene expression). Once mapped, those QTLs can be colocalized with GWAS risk variants to identify putative causal mechanisms. We and others have colocalized novel or existing expression QTL data sets with OA GWAS to identify putative causal genes (would add refs for we and others). Despite these successes, many OA GWAS loci remain unexplained. One possible reason is that the disease associated variants might influence traits other than gene expression. This may be due to alternative splicing as existing literature has described substantial contributions of both expression QTLs and splicing QTLs to disease risk; however, these disease-associated eQTLs and sQTLs often influence disease risk via different genes^7–9^. While OA tissue is characterized by distinct changes in splicing patterns^10^; sQTLs have yet to be mapped in cartilage or other OA-relevant cell types.

To address this critical need, we mapped splicing QTLs in primary human chondrocytes, the primary cell type present in articular cartilage. Splicing QTLs were mapped in chondrocytes both in resting state and in response to a fibronectin fragment (FN-f), a known OA trigger. FN-f is present in the cartilage and synovial fluid of OA patients^11^ and in cell culture studies has been shown initiate a range of catabolic signaling pathways characteristic of chondrocytes isolated from OA tissue, mimicking the OA-like state^12–14^. This approach allows us to capture the transition from a normal to an OA phenotype and identify variants that exert effects in both homeostatic and pathological settings.

We analyzed RNA-seq data from 101 human chondrocyte samples treated with either PBS (control) or FN-f and 16 OA donor samples. We identified hundreds of genes that were differentially spliced between PBS-treated and either FN-f treated or OA chondrocytes. Differential splicing events induced by FN-f mimicked those seen in OA tissue underscoring the accuracy of our FN-f model. We used CRISPR to mimic a splicing event observed in both FN-f treated normal chondrocytes and in OA chondrocytes and found that it induced OA-like expression changes. We identified 7,188 splicing quantitative trait loci (sQTL) affecting 3,056 genes, over 1,000 of which exhibited condition-specific effects. Colocalization of these sQTLs with OA GWAS identified 6 putative OA risk genes, including one novel candidate, *PBRM1*. This approach demonstrates the potential of sQTL analysis to uncover novel genetic factors contributing to OA susceptibility. Our findings both improve the mechanistic understanding of the disease and identify genes that can be further studied and potentially targeted for therapeutic intervention.

## Results

### FN-f induces alternative splicing events in human chondrocytes

To determine how FN-f stimulation influences RNA splicing we reanalyzed our existing RNA-seq data sets obtained from primary human chondrocytes isolated from 101 non-OA donors treated with either PBS or FN-f, a known OA trigger. Initial quality control revealed minor batch effects associated with the RNA extraction kit and FN-f batch (**Fig. S1a**). We corrected these technical confounders using the limma package^15^, resulting in improved sample clustering by biological condition (**Fig. S1b**). Differential analysis using leafcutter identified 974 differential splice junctions corresponding to 590 genes (adjusted p < 0.05, |ΔPSI| > 0.15, **Table S2**). Hierarchical clustering of differentially spliced intron junctions revealed distinct patterns between PBS control and FN-f treated samples (**Fig. 1a**). Over ninety percent of these splicing events were previously annotated which supports the validity of our splicing analysis (**Fig. 1b**). We performed pathway and Gene Ontology (GO) enrichment analysis on the differentially spliced genes to understand the potential functional implications of these splicing changes. For FN-f-stimulated chondrocytes, this analysis revealed significant enrichment in several pathways and processes some of which had possible connections to OA pathogenesis (**Supplementary Fig. 2a**). Key enriched pathways included signal transduction, FGFR1 signaling in disease, DCC (deleted in colorectal cancer)-mediated attractive signaling, and osteoclast differentiation. Enriched GO terms included anatomical structure morphogenesis, actin cytoskeleton organization, and cell motility.

**Fig 1.**
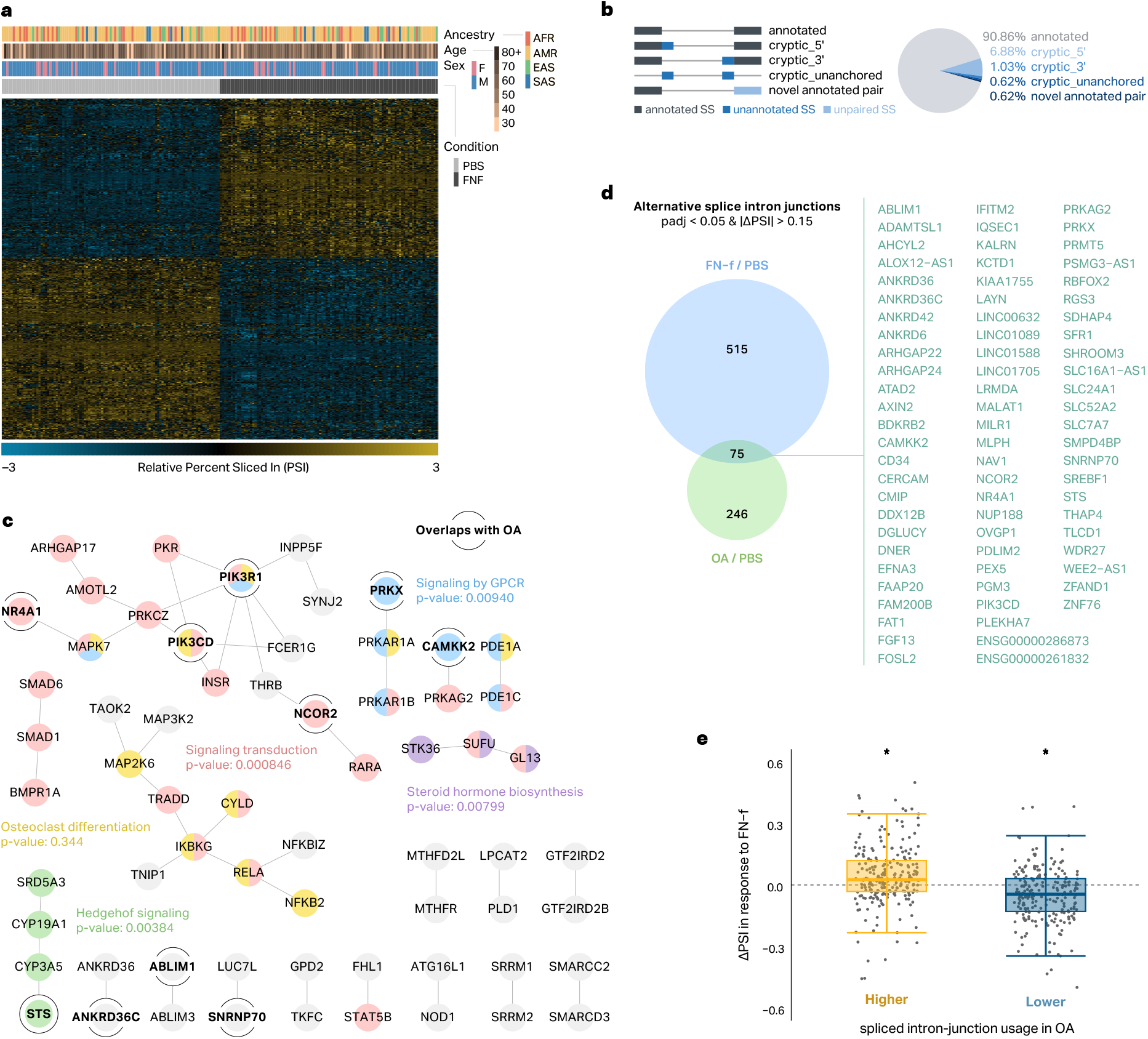
FN-f induces alternative splicing events in human chondrocytes. (**a**) a A heatmap of differential splicing events between PBS- and FN-f-treated samples. (**b**) A pie chart illustrates the distribution of differential alternative splicing intron junction types. (**c**) Protein-protein interaction network of differentially spliced genes using high-confidence interactions (Interaction score >0.900). Colors highlight membership in selected pathways. Yellow circles indicate genes that are also differentially spliced in OA tissue. (**d**) A Venn diagram depicts the overlap between FN-f and OA-associated splicing changes. (**e**) Boxplots depict the log 2 fold-change in PSI in response to FN-f for introns that were differentially spliced in OA chondrocytes. * indicates Wilcoxon test p-value < 0.01.

To further interrogate the relationship between differentially spliced genes we performed protein-protein interaction network analysis and detected 64 nodes with high-confidence interactions (Interaction score > 0.900). This network highlighted subnetworks corresponding to critical pathways such as GPCR signaling, signal transduction, osteoclast differentiation, hedgehog signaling, and steroid hormone biosynthesis (**Fig. 1c, Fig. S2a**). Taken together these results demonstrate that FN-f induces large-scale changes in alternative splicing that converge on specific pathways and processes, many of which have possible relevance to OA.

### FN-f-treated chondrocytes and chondrocytes from OA tissue exhibit similar splicing patterns

To determine if our FN-f-treated chondrocytes exhibited similar splicing patterns to those found in OA tissue we performed RNA-seq on chondrocytes isolated from cartilage obtained from 16 donors who underwent knee replacement surgery. Comparison between OA chondrocytes and our PBS treated non-OA chondrocytes revealed 466 significantly differentially spliced intron junctions corresponding to 321 genes(adjusted p < 0.05, |ΔPSI| > 0.15, **Fig. S2b, Table S2**). Genes that were differentially spliced in OA compared to non-OA chondrocytes (treated with PBS) were enriched in multiple KEGG pathways including MCM complex, Rho GTPase cycle, and fatty acid metabolism (**Fig. S2d, Table S3**) and GO terms including positive regulation of GTPase activity, cell morphogenesis, and regulation of filopodium assembly. 33 intron junctions (corresponding to 33 distinct genes) were significantly altered in both FN-f-stimulated and OA chondrocytes (**Fig. 1d**, right panel). Several of these OA-associated differential spliced genes overlapped the protein-protein interaction nodes that we identified from FN-f induced splicing events (**Fig. 1c** in yellow). These genes included several that have been associated with cartilage biology or processes relevant to OA pathogenesis. For instance, *SNRNP70* is splicing factor that has been implicated in rheumatiod arthritis^16^; *MALAT1* is a lncRNA involved in inflammation and chondrocyte proliferation^17^, and *NCOR2* is a nuclear receptor co-repressor that is hypermethylated and down-regulated in OA tissue^18^.

Next, we sought to determine the extent to which splicing events in FN-f treated chondrocytes mirror those seen in OA. We divided the 466 introns that were differentially spliced between OA and non-OA (PBS-treated) chondrocytes into those that showed increased or decreased percent spliced in (PSI) in OA samples. Then we plotted the log 2 fold-change of PSI in FN-f vs PBS treated chondrocytes. On average, introns that exhibited increased PSI in OA tissue also exhibited increased PSI in response to FN-f (**Fig 1e**, Wilcox test p-value < 0.01). Conversely, introns that exhibited decreased PSI in OA tissue also exhibited decreased PSI in response to FN-f (**Fig 1e**, Wilcox test p-value < 0.01). The agreement between splicing changes in OA and FN-f-treated chondrocytes, both in terms of broad directional changes and specific high-confidence alterations, suggests that the FN-f-treated cells are a valuable model to interrogate OA-related alternative splicing events. This similarity further validates the FN-f stimulation model for studying OA-related splicing dysregulation and provides valuable insights into the molecular mechanisms potentially underlying OA pathogenesis. The identified set of consistently altered splicing events offers a focused group of targets for future studies to understand and potentially intervene in OA progression.

### SNRNP70 alternative exon 8 deletion induces OA-like expression patterns

We identified *SNRNP70*, a critical component of the U1 snRNP complex, as differentially spliced in both FN-f-stimulated and OA chondrocytes compared to PBS-treated controls. RNA-seq analysis revealed significant changes in *SNRNP70* splicing, particularly between exons 7 and 8 (**Fig. 2a**). The alternative exon 8 (Chr19:49,102,114-49,103,587) showed reduced inclusion in FN-f and OA conditions, with delta PSI differences of 22% (FN-f/PBS) and 23% (OA/PBS). This indicates alternative exon 8 skipping in 67% and 68% of transcripts in FN-f and OA conditions, respectively, compared to 45% in PBS controls. Exon-level expression analysis confirmed significant downregulation (p < 0.01) of the alternative exon 8 in both conditions (**Fig. 2b**).

**Fig 2.**
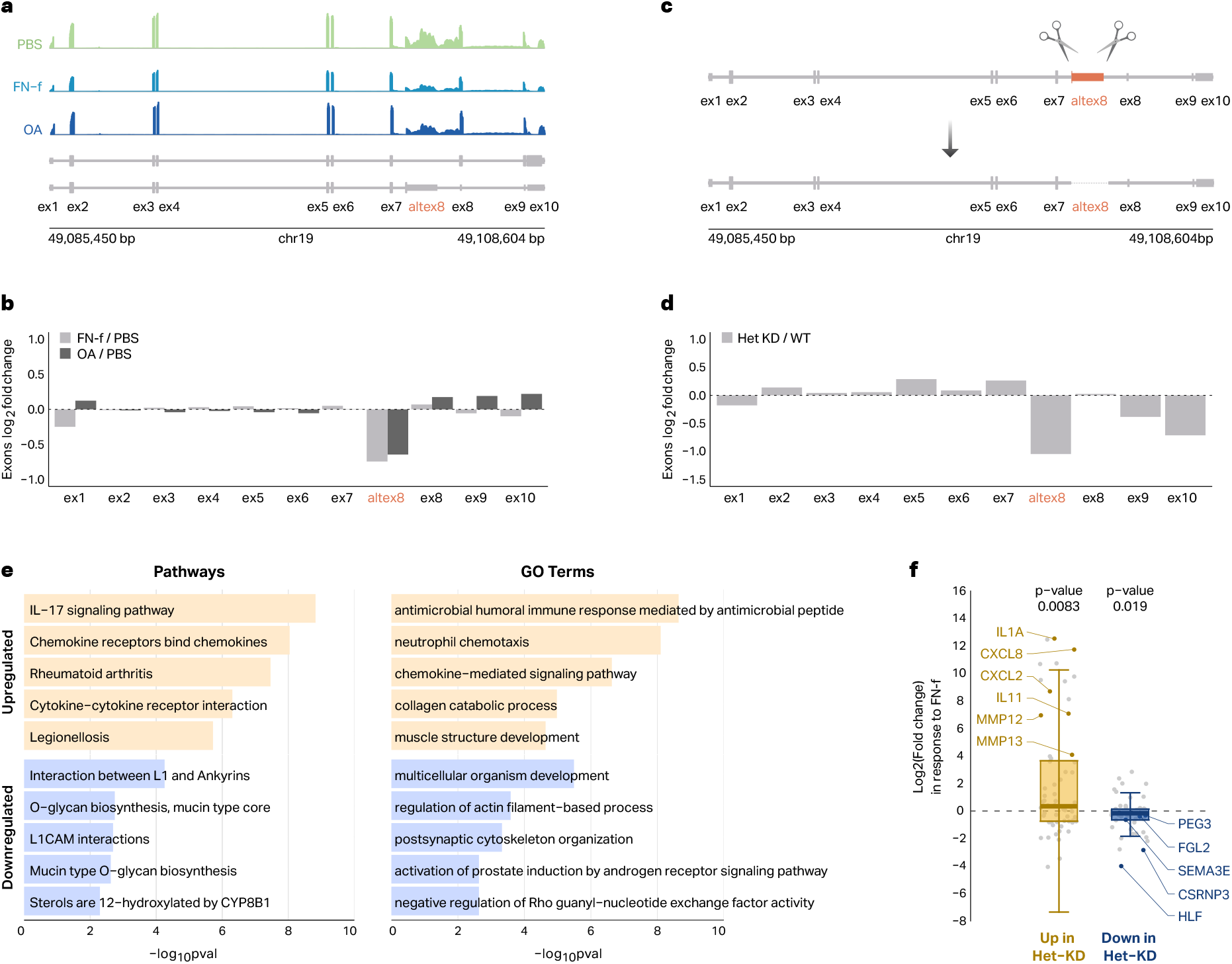
Deletion of *SNRNP70* alternative exon 8 mimics OA-related splicing patterns. (**a**) RNA-seq signal track for PBS-treated, FN-f-treated, and OA chondrocytes at the *SNRNP70* gene locus. Below the signal tracks are transcript annotations representing the MANE Select transcript (ENST00000598441.6, top), and a transcript containing the alternative exon 8 (ENST00000401730.5, bottom). (**b**) Barplot comparing log2 fold change for expressed exons with Fn-f (gray bars) and OA (black bars) as compared to PBS. (**c**) Schematic of CRISPR/Cas9 genome editing strategy for *SNRNP70* alternative exon 8 deletion. (**d**) Barplot comparing log2 fold change for expressed exons after *SNRNP70* alt ex8 deletion. (**e**) Barplot of KEGG & Reactome pathways and GO terms enriched in differentially expressed genes between alt ex8 heterozygous knockdown (Het-KD) and wildtype (WT). (**f**) Boxplot showing genes that are up- or downregulated in alt ex8 het-KD vs. controls exhibit the same directions in FN-f stimulated chondrocytes compared to PBS control. Right (Yellow): Highlighted examples of known OA-related genes upregulated. Left (Blue): Highlighted examples of known OA-related genes downregulated.

To assess the functional role of *SNRNP70* alternative exon 8, we employed CRISPR-Cas9 to delete this region in primary human chondrocytes (**Fig. 2c**). Our deletion strategy targeted guide RNAs to both sides of a 1,735 bp region encompassing the alternative exon 8 (**Supplementary Fig. 3a**). PCR screening of 46 single-cell-derived colonies revealed 26% with heterozygous deletion of the alternative exon 8 (**Supplementary Fig. 3b,c**). None of the colonies exhibited homozygous deletion which could be related to *SNRNP70’s* classification as an essential gene^19^. Sanger sequencing of Het-KD colonies confirmed the deletion, with an additional cytosine deletion beyond the expected cut site (**Supplementary Fig. 3d**).

To determine the phenotypic impact of exon 8 deletion we performed RNA-seq on edited and unedited colony-expanded cells. The differential analysis confirmed decreased *SNRNP70* alternative exon 8 inclusion (**Fig. 2c**) and identified 135 differentially expressed genes (DESeq2, FDR-adjusted p-value < 0.01). Genes that were upregulated in Het-KD cells were enriched for multiple GO terms and KEGG pathways including IL-17 signaling, chemokine receptor binding, and rheumatoid arthritis (**Fig. 2e**, Supplementary Data 4). Genes that were downregulated in Het-KD cells were enriched for pathways including L1 and Ankyrin interactions and O-glycan biosynthesis (**Fig. 2e**, Supplementary Data 4).

Interestingly, the directionality of gene expression changes in Het-KD cells significantly correlated with those in FN-f-stimulated chondrocytes (p < 0.05 for both up-and down-regulated genes, **Fig. 2f**). Genes that were upregulated both in response to FN-f and in response to alternative exon 8 deletion included *CXCL8, CXCL2, IL11, MMP12*, and *MMP13*, which are associated with inflammation and extra-cellular matrix degradation in OA^20–22^. Downregulated genes included *PEG3, FGL2, SEMA3E, CSRNP3*, and *HLF*, which have been implicated in chondrocyte homeostasis^23–27^. These results suggest that the exclusion of *SNRNP70* alternative exon 8 in response to cartilage matrix damage modeled by FN-f treatment may be responsible for some of the other expression changes associated with the OA phenotype.

### Genetic variants impact splicing in resting and activated chondrocytes

We performed splicing QTL (sQTL) analy-sis on both PBS control and FN-f stimulated chondrocyte samples to determine the impact of genetic differences on chondrocyte mRNA splicing. We tested the association of each RNA splicing event with genetic variants within ±100 kb of the start and end points of the splice intron junctions, using PSI values adjusted for confounding factors (**Supplementary Fig. 4a)**. We identified a total of 5,873 unique sQTLs corresponding to 2,575 sGenes (genes with at least one significant sQTL) in PBS control, and 4,606 sQTLs corresponding to 2,132 sGenes in FN-f stimulated chondrocytes (QTLtools^28^, FDR < 0.05, **Fig. 3a, Supplementary Data 5**).

**Fig 3.**
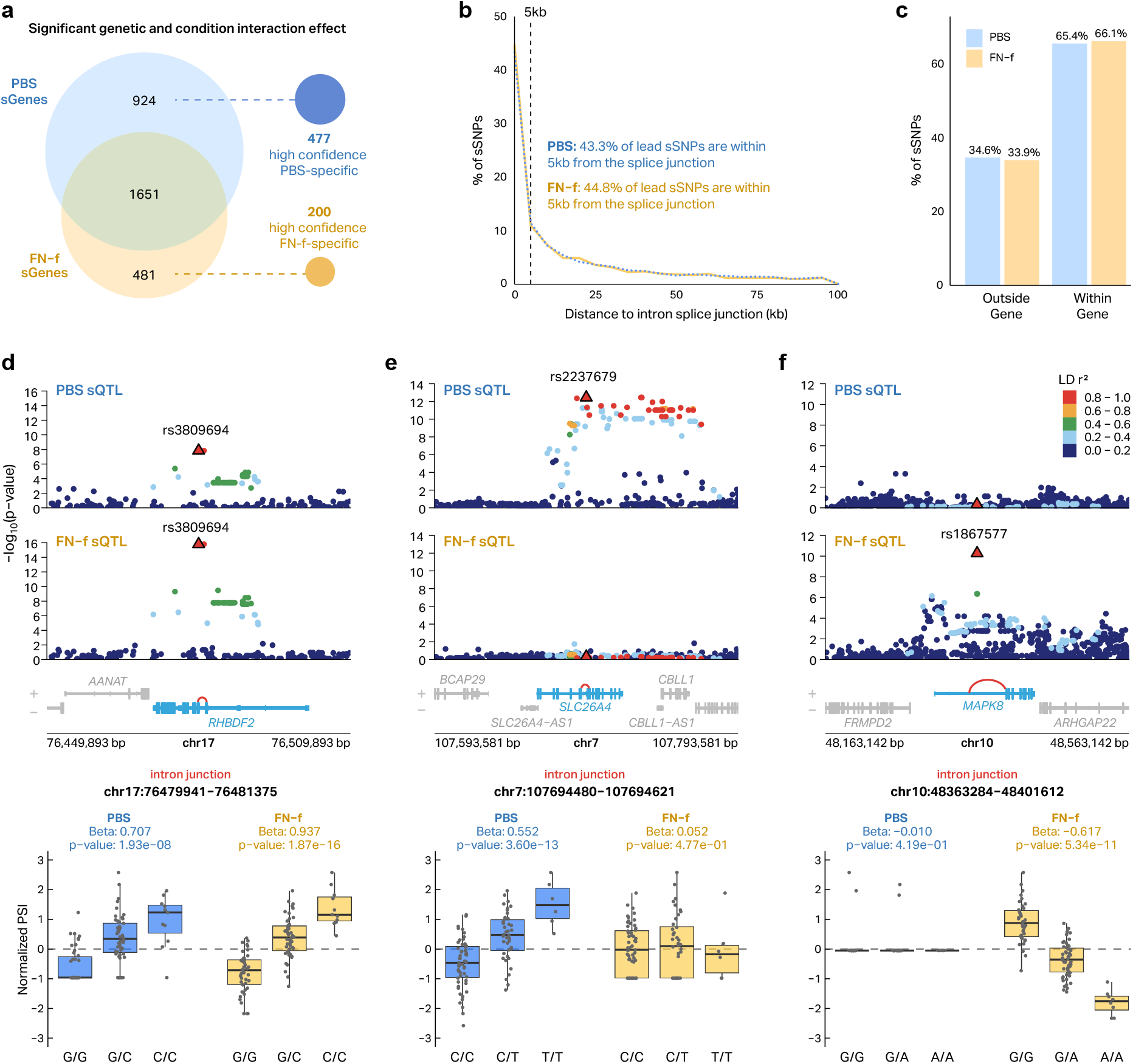
Condition-specific splicing quantitative trait loci (sQTL) discovery in resting and activated chondrocytes. (**a**) Venn diagram showing the overlap between sGenes identified in PBS and FN-f treated conditions. Right: Linear mixed-effect modeling allowed for the detection of 677 condition-specific sGenes (**b**) The distribution of distances between sSNPs and the affected splice intron junctions (kb). (**c**) Percentage of sSNPs located within the same gene (within a gene) or outside of the gene. Examples of (**d**) shared, (**e**) PBS-specific, and FN-f-specific (**f**) sQTLs.

Conditional analysis revealed 182 secondary signals, 13 tertiary signals, and 2 quaternary signals for PBS-treated chondrocytes and 121 secondary signals and 1 tertiary signal for FN-f-treated chondrocytes (**Supplementary Fig. 4b**). In total, 116 PBS sGenes and 82 FN-f sGenes had two or more independent signals. For example, the *CAST* gene exhibited multiple independent signals associated with the splice intron junction chr5:96768431-96776404 in PBS control, while the *MICA* gene exhibited multiple independent signals with the splice intron junction chr6:31410797-31411209 in FN-f stimulated chondrocytes (**Supplementary Fig. 5**).

We analyzed the genomic distribution of sQTLs (sSNPs) relative to their associated splice intron junctions. Lead sSNPs were largely close to splice junctions with 43% and 45% of lead SNPs within 5Kb of the affected spice junctions for PBS and FN-f respectively (**Fig. 3b**). Further analysis revealed that the majority of sSNPs were located within the same gene as their associated splice intron junction **(Fig. 3c)**. Taken together these results both support the validity of our sQTLs and suggest that sSNPs largely act at the affected splice junctions rather than via distal mechanisms.

To determine if any of the effects of variants were dependent on the condition, we used a linear mixed-effect model and identified 1635 sGenes that exhibited significant genetic and condition interaction effects (lme4^29^, ANOVA, p < 0.05). To compile a high-confidence, large-effect size list of condition-specific sQTLs, we further refined this list to include only those that were identified as an sQTL in only one condition and exhibited an absolute beta difference between conditions of greater than or equal to 0.2. In total, we identified 677 high-confidence condition-specific sGenes (**Fig 3a**). 400 lead SNPs had a stronger genetic effect on splicing in PBS-treated cells and 277 lead SNPs had a stronger genetic effect on splicing in FN-f-treated cells. Examples of shared, PBS-specific, and FN-f-specific sQTLs are depicted in **Figure 3d-f**. Several of these condition-specific sGenes have known roles in cartilage biology and/or OA. For example, **Fig. 3e** shows a PBS-specific condition sQTL for *SLC26A4* involving the splice junction chr7:107694480-107694621. SLC26A4, an anion transporter, may influence chondrocyte metabolism in homeostatic conditions, potentially affecting cartilage maintenance^30^. **Fig. 3f** demonstrates an FN-f-specific sQTL for *MAPK8*, associated with the splice junction chr10:48363284-48401612. MAPK8 (JNK1) is involved in inflammatory signaling. MAP kinases are known to play a key role in the regulation of matrix-degrading metalloproteases and their inhibition is currently being investigated for therapeutic treatment of OA^31,32^.

### RNA-binding proteins provide insight into putative sQTL mechanisms

RNA-binding proteins (RBPs) play crucial roles in post-transcriptional regulation, influencing various cellular processes, including inflammation and metabolic changes related to OA^33,34^. Recent studies have highlighted the dysregulation of RBPs in chondrocytes, synovial fibroblasts, and osteoblasts, suggesting their significant impact on OA progression^35,36^.

To gain insight into the putative mechanisms through which our sQTLs were influencing splicing in chondrocytes, we leveraged publicly available enhanced crosslinking and immunoprecipitation (eCLIP) data from the ENCODE database^37^. This dataset encompasses binding sites for 140 RBPs across three cell types: HepG2 (n = 85), K562 (n = 118), and SM-9MVZL (n = 2). While these cell lines are not OA-specific, similar data is not currently available in chondrocytes and presumably many of these binding sites are shared across cell types.

Using the QTLtools fenrich module, we assessed the statistical significance of RBP enrichment through permutation tests. For PBS control lead sQTLs, we identified 12 significantly enriched RBPs (odds ratio > 1, empirical P < 0.05), including SF3B4, EFTUD2, and PRPF8. The FN-f condition yielded 7 enriched RBPs, including UP3, EFTUD2, and SF3B4 (**Fig. 4a**, Supplementary Data 6). We found that 255 sQTLs associated with intron junctions were enriched for at least one RBP and showed a significant correlation with RBP gene expression (Pearson’s R^2^ > 0.15, P < 0.01; **Supplementary Data 6**).

**Fig 4.**
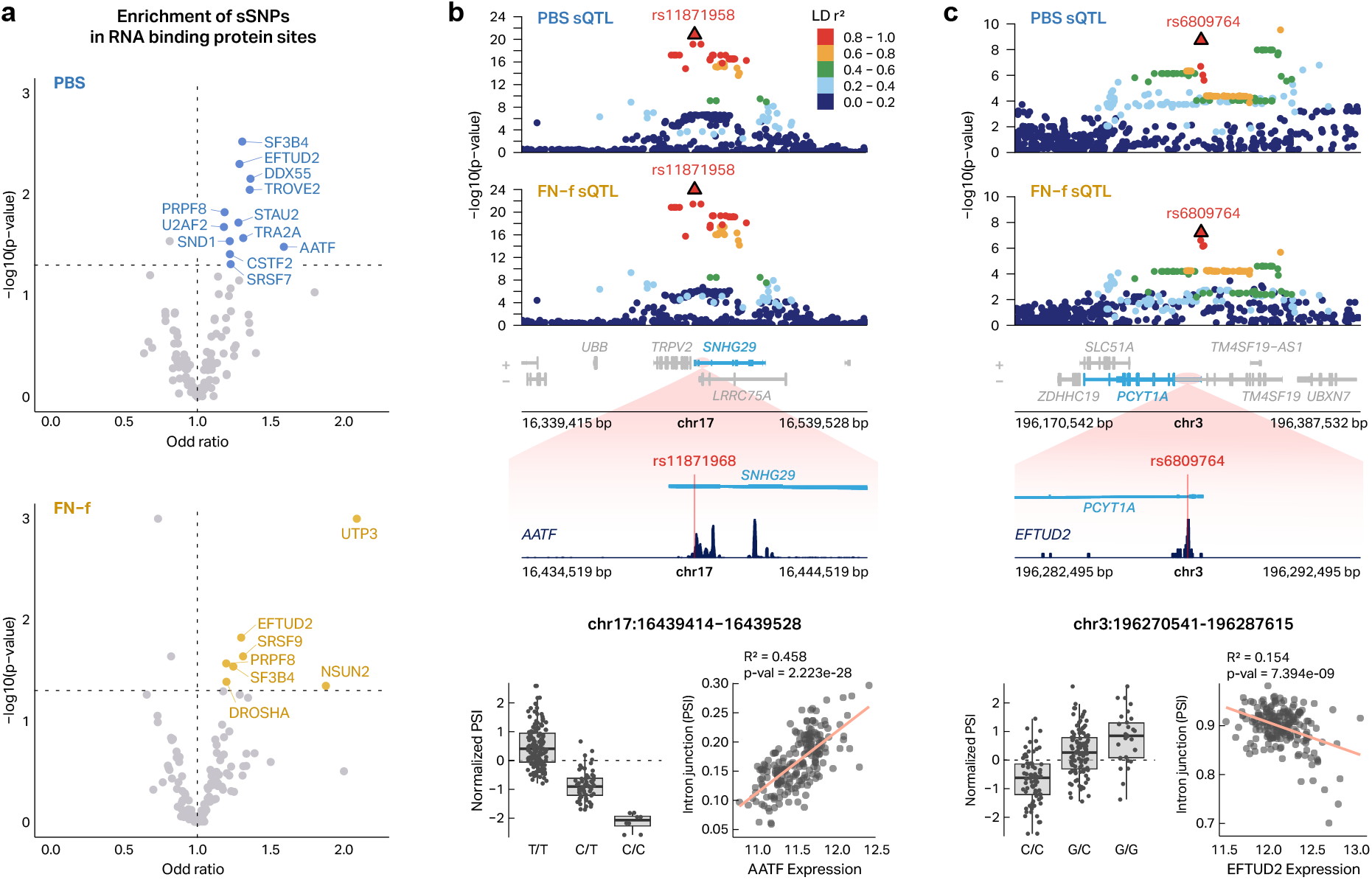
RNA-Binding protein associations with sQTLs. (**a**) Volcano plot depicting the effect size (x-axis) and -log10 p-values (y-axis) of RBP sites enrichment in sQTLs identified in PBS- or FN-f-treated chondrocytes. Data points are colored blue (PBS control) or orange (FN-f) for p-value < 0.05 and odds ratios > 1. (**b**)A locus zoom plot for an sQTL in the *SNHG29* gene (top) with a zoom-in showing an AATF binding site overlapping the sSNP rs11871958 (middle). (Bottom) Looking across donors AATF expression is correlated with PSI of this intron lending support for its role in regulating this splicing event. (**c**) Example of *PCYT1A* gene and rs6809764 sQTL shared region with EFTUD2 RBP. This panel follows the same format as panel b.

An example of a putative RBP-meditated sQTL is rs11871958 which is associated with alternative splicing of *SNHG29* and overlaps an AATF binding site (**Fig. 4b**). The minor C allele of rs11871958 is associated with decreased PSI of the associated intron junction (chr17:16439414-16439528). Interestingly, *AATF* gene expression is strongly correlated with the PSI of the associated intron junction lending further support to the role of AATF in this splicing event (R^2^ = 0.458, P = 2.223e-28). *SNHG29*, a long non-coding RNA, has been previously reported to regulate the miR-223-3p/CTNND1 axis and influence the Wnt/β-catenin signaling pathway in glioblastoma^38^ which has been previously implicated in OA pathogenesis^39^.

Another example is rs6809764 which was identified as an sQTL for *PCYT1A* and overlaps an EFTUD2 binding site (**Fig. 4c**). The minor G allele of rs6809764 is associated with increased PSI of the intron junction (chr3:196270541-196287615). Expression of *EFTUD2* correlated with decreased junction usage (R^2^ = 0.154, P = 7.394e-09) which supports EFTUD2 as a possible mediator of this sQTL. *PCYT1A* encodes an enzyme crucial for phosphatidylcholine biosynthesis and has been observed to localize to the nuclear envelope in hypertrophic chondrocytes of bone growth plates, coinciding with a significant increase in cell volume^40^. This localization suggests a potential role for *PCYT1A* in chondrocyte morphology changes during hypertrophy, a process relevant to cartilage development and potentially to OA pathogenesis.

These findings provide insights into the regulatory mechanisms of OA-associated splicing events and highlight potential new targets for therapeutic intervention. However, we acknowledge the limitations of using non-OA-specific cell lines for RBP binding data. Further biochemical studies in relevant cell types will be crucial to fully elucidate the regulatory programs orchestrating disease-related splicing changes in OA.

### Colocalization analysis reveals putative OA risk genes

To understand the potential role of splicing in OA risk, we performed colocalization analysis between our sQTLs and 100 independent OA GWAS loci spanning 11 OA-related phenotypes reported by Boer et al^4^. We selected sQTLs whose lead SNP was in high LD (r^2^ > 0.5) with a lead SNP from any of the 11 OA GWAS data SNPs and tested for colocalization using the *coloc* package(**Supplementary Data 7**). In total, we identified 6 colocalized signals corresponding to 6 unique sGenes across 4 OA phenotype subtypes (**Table 1, Supplementary Data 7**). For *GCAT*, we identified sQTLs in both conditions (rs13057823 in PBS and rs2071910 in FN-f) that colocalize with the Total Hip Replacement (THR) GWAS signal (index variant rs12160491). Both sQTLs affect the same splice intron junction (chr22:37808163-37809949), suggesting a consistent impact on GCAT splicing across conditions (**Fig. 5b**). The *GCAT* locus demonstrated strong evidence of colocalization in both conditions, with PP H4 / (PP H3 + PP H4) ratios of 0.9392 in PBS and 0.8955 in FN-f. The *HMGN1* locus revealed a PBS-specific sQTL (rs2249666) colocalizing with a total joint replacement (TJR) GWAS signal (index variant rs9981884). This sQTL affects the splice intron junction chr21:39347292-39347379, with a colocalization probability (PP H4 / (PP H3 + PP H4)) of 0.8877 (**Fig. 5c**). The sQTL rs1886248 for *RNF144B* colocalized with the FingerOA GWAS signal (index variant rs9396861) in both PBS and FN-f conditions (**Fig. 5d**). The *WWP2* locus showed PBS-specific colocalization, with sQTL rs7192245 colocalizing with the KneeOA GWAS signal (index variant rs34195470) (**Fig. 5e**). The colocalization probability (PP H4 / (PP H3 + PP H4)) for this locus was 0.7136. For *COLGALT2*, we identified a PBS-specific sQTL (rs74767794) that colocalizes with a THR GWAS signal (index variant rs1327123) (**Fig. 5f**). This locus demonstrated a colocalization probability (PP H4 / (PP H3 + PP H4)) of 0.9683. For the *PBRM1* signal, we identified an FN-f-specific sQTL (rs7628578) colocalizing with a THR GWAS signal (index variant rs3774354). This sQTL influences the splice intron junction chr3:52658315-52662133 (**Fig. 5a**), with moderate evidence of colocalization (PP H4 / (PP H3 + PP H4) = 0.7263) in the FN-f condition.

**Table 1.**
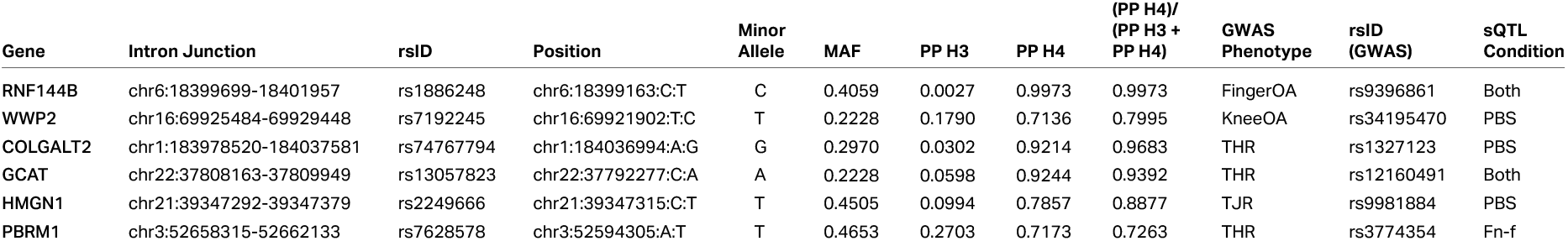
Chondrocyte sQTLs that colocalize with OA GWAS loci. The table shows the significant results of colocalization between sQTL and GWAS signals (PP H4 > 0.7) with the most significant splice intron junction. (PP, value of posterior probability; FingerOA, OA of the finger; KneeOA, OA of the knee; THR, Total Hip Replacement).

**Fig 5.**
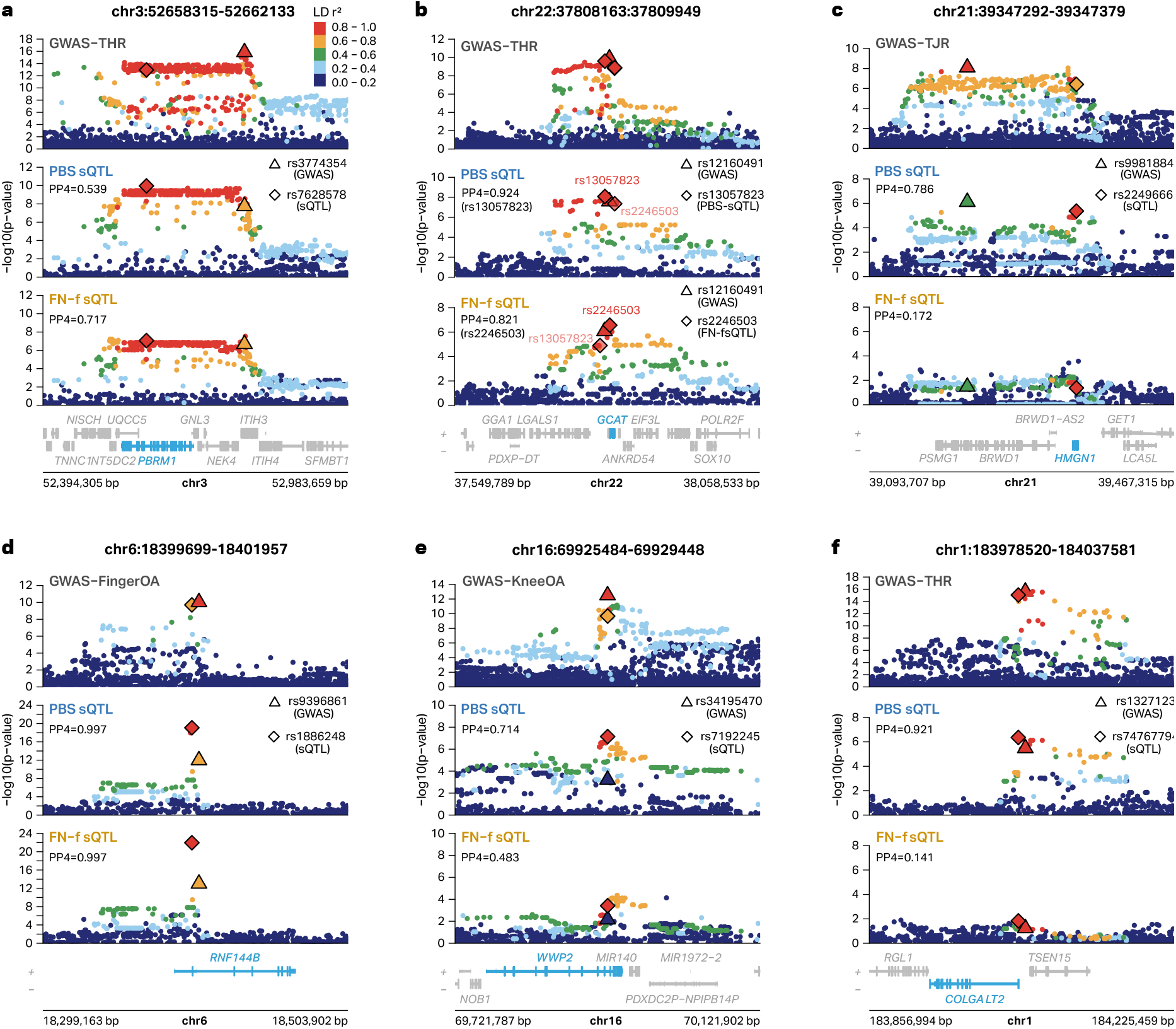
Six sQTLs colocalize with OA GWAS signals. (**a**) A THR GWAS signal cololocalizes with an sQTL found in both conditions for *PBRM1* associated with splice intron junction chr3:52658315-52662133. (**b**) THR GWAS signal cololocalizes with an sQTL found in both conditions for *GCAT* associated with splice intron junction chr22:37808163-37809949. (**c**) A TJR GWAS signal cololocalizes with an sQTL found in both PBS-treated chndrocytes for *HMGN1* associated with splice intron junction chr21:39347292-39347379. (**d**) A Finger OA GWAS signal cololocalizes with an sQTL found in both conditions for *RNF144B* associated with splice intron junction chr6:18399699-18401957. (**e**) A Knee OA GWAS signal cololocalizes with an sQTL found in PBS-treated chondrocytes for *WWP2* associated with splice intron junction chr16:69925484-69929448. (**f**) A THR GWAS signal cololocalizes with an sQTL found in PBS-treated for *COLGALT2* associated with splice intron junction chr1:183978520-184037581.

To the best of our knowledge, this is the first time any of these splicing events have been colocalized with OA GWAS signals; however, three of these genes have been implicated in OA via colocalization with expression QTLs. *GCAT* and *HMGN1* have been previously associated with knee and/or hip OA via colocalization with an osteoclast-specific eQTL^41^. *RNF144B* was previously linked to OA via colocalization between OA GWAS variants and eQTLs from chondrocytes, tibial nerves, and testis^4,42^. Our results build upon these previous studies by providing more mechanistic detail into how these variants act, presumably via their impact on alternative splicing

Three of our colocalized genes have not been previously linked to OA via colocalized eQTLs. *WWP2* and *COLGALT2* have been implicated in OA by their proximity to mQTLs that colocalize with OA GWAS variants and control methylation sites near WWP2 and COGALT2^43^. These associations were then confirmed via epigenome editing^44,45^. To the best of our knowledge, one of our colocalized sGenes (*PBRM1*) has not been previously linked to OA via genetic methods. These findings highlight the potential role of alternative splicing in mediating OA risk and demonstrate the value of integrating sQTL data with GWAS results to uncover functional mechanisms underlying genetic associations.

## Discussion

To our knowledge, this study is the first comprehensive sQTL map in chondrocytes to date, providing new insights into the relationship between genetic variation, alternative splicing, and OA risk. By integrating sQTL analysis with OA GWAS data and functional validation, we have identified potential novel mechanisms underlying OA pathogenesis and highlighted putative OA risk genes for further investigation in future therapeutic development.

The splicing alteration observed in both FN-f stimulated and OA chondrocytes highlighted the role of alternative splicing in OA progression. Our identification of 974 and 466 differentially spliced intron junctions in FN-f-stimulated and OA chondrocytes, respectively, suggests broad splicing dysregulation in OA. The overlap between FN-f-induced and OA-associated splicing events supports the relevance of our ex vivo model and suggests that some of these splicing changes may contribute to OA pathogenesis.

Our functional interrogation of *SNRNP70* alternative splicing provides evidence for the impact of splicing changes on OA-related gene expression. The heterozygous deletion of *SNRNP70* alternative exon 8 was a mimic of the reduced retention of alternative exon 8 that occurs during OA and in response to FN-f. This deletion led to alterations in genes and pathways crucial for OA progression, including inflammatory signaling and matrix degradation, and suggests that alternative spliced genes may influence cellular phenotypes relevant to OA. The identification of other differentially spliced genes in both FN-f stimulated and OA chondrocytes, such as *STS, ANKRD36C*, and *PIK3CD*, provide potential targets for further investigation in OA.

Our sQTL analysis identified genetic variants influencing splicing in both resting and activated chondrocytes. The prevalence of local regulatory effects suggests that cis-acting splicing regulation contributes to how genetic variation influences OA risk. The colocalization analysis between our sQTLs and OA GWAS signals identified 6 colocalized signals. Three of these colocalized genes have been previously linked to OA via colocalized eQTLs but our results build on this knowledge by suggesting that their pathogenic mechanism may act via alterations to mRNA splicing. While *WWP2* has been functionally linked to OA through knockout studies, both *WWP2* and *COLGALT2* are implicated in OA by their proximity to colocalized mQTLs, despite the absence of direct eQTL associations^43^.

One of our colocalized genes *PBRM1* is novel and provides a new potential OA mediator for further research and potential therapeutic development. The rs3774354 GWAS signal was initially assigned to ITIH1 based on genomic proximity^4^, however, our colocalized sQTL suggests that *PBRM1* may be the causal gene at this locus^4^. This suggests that altered splicing of PBRM1, rather than effects on ITIH1, may be the mechanism driving OA risk at this locus. As a SWI/SNF chromatin-remodeling complex component^46^, *PBRM1* influences gene expression patterns. Its role in mesenchymal stromal cell differentiation is particularly intriguing in the context of OA. Previous research has shown that *PBRM1* affects osteogenic differentiation and modulates *SOX9* expression, a key regulator of chondrogenesis and expression of chondrocyte matrix genes^47,48^.

These observations suggest that *PBRM1* might be involved in maintaining the delicate balance between different cell fates in joint tissues, a process often disrupted in OA.

Interestingly, *GCAT* and *HMGN1* have been associated with hipOA, knee and/or hip OA and/or allOA in osteoclast-specific eQTL^41^. Our findings extend this by providing evidence for their involvement in chondrocyte splicing regulation. *GCAT*, encoding glycine C-acetyltransferase, plays a crucial role in glycine metabolism in mitochondria^49^. Previous studies have shown that epigenetic downregulation of *GCAT* can lead to respiration defects in aged fibroblasts, and glycine supplementation was found to restore these defects^49^. While the direct link between GCAT and OA pathogenesis remains to be fully elucidated, our identification of GCAT through sQTL analysis suggests that alterations in mitochondrial function and glycine metabolism may contribute to OA risk, possibly through effects on cellular energy production or extracellular matrix synthesis. *HMGN1* is a known nonhistone chromatin remodeler, and has been implicated in regulation of chondrogenesis^50^. Furusawa et al demonstrated *HMGN1* directly regulated expression level of the SOX9 gene. This suggests that alternative splicing may modulate *HMGN1* function and regulate chondrocyte-specific gene expression.

*RNF144B* showed different patterns of condition specificity in our sQTL analysis compared to previous eQTL studies. We observed strong *RNF144B* sQTLs in both PBS- and FN-f-treated conditions, whereas the eQTL for *RNF144B* was highly specific to the FN-f-treated condition. This discrepancy highlights how sQTL and eQTL analyses can provide complementary information for mRNA production and processing. Given *RNF144B*’s role in regulating DNA damage-induced apoptosis, its differential splicing could affect chondrocyte survival and turnover in OA.

*WWP2* has been previously implicated in OA through methylation studies. In particular, Shepherd et al. identified a differentially methylated region (DMR) associated with the OA risk-conferring allele of rs34195470^44^. This DMR was found to act as a methylation-sensitive transcriptional repressor, with increased DNA methylation resulting in increased *WWP2* expression, particularly of the WWP2-FL and WWP2-N transcripts. Our colocalization of a *WWP2* sQTL and OA GWAS signal provides a direct tie to OA and adds another layer to its regulatory complexity in OA, perhaps suggesting that OA risk variants alter splicing via changes to DNA methylation.Further work is necessary to explore this hypothesis.

*COLGALT2* has also been previously associated with OA risk^45,51^. Previous research has identified that the OA risk allele of rs11583641 corresponded to reduced DNA methylation at nearby CpGs and increased *COLGALT2* expression^52^. Our sQTL findings now implicate alternative splicing as another potential mechanism by which *COLGALT2* variation may influence OA risk, further highlighting the complex regulatory landscape of this gene in OA pathogenesis.

While our study represents a significant advance in understanding the role of splicing in OA, it has several limitations that should be addressed in future research. Our sample size of 101 donors, while substantial for an sQTL study in chondrocytes, may not have captured all splicing-related OA GWAS genes. Future studies with larger sample sizes will be crucial to fully elucidate the splicing landscape in OA. Additionally, while chondrocytes are highly relevant to OA, investigating splicing changes in other cell types involved in joint biology will provide a more comprehensive understanding of OA pathogenesis. Our FN-f stimulation model, while valuable, cannot fully recapitulate the complex joint environment. Advanced 3D culture systems, such as cartilage organoids or joint-on-a-chip models, could provide more physiologically relevant contexts for validating our findings. Furthermore, our study focused on a single time point of FN-f stimulation. Future studies considering other time-points may capture the splicing changes not present here. This could involve analyzing samples at multiple time points after FN-f stimulation or, in the case of clinical samples, collecting data from patients at different stages of OA progression. The genetic background of our cohort, while diverse, may not capture all population-specific effects on splicing in OA. Future studies including more diverse populations could identify potential population-specific splicing QTLs and their relevance to OA risk and progression.

While our functional validation of *SNRNP70* provides valuable insights, further experimental work is needed to understand the functional consequences of the colocalized sQTLs we identified. Elucidating the impact of these splicing changes on OA pathogenesis remains challenging and will require extensive investigation. Future studies could employ a range of approaches to validate and characterize the effects of colocalized sQTLs. Particular focus should be given to the novel gene *PBRM1*, as well as *GCAT* and *HMGN1*, which were previously identified in osteoclast eQTL colocalized with OA GWAS signals but now show evidence of splicing regulation in chondrocytes. Additionally, further investigation into *WWP2* and *COLGALT2*, which have been previously implicated in OA through other mechanisms, could provide insights into the interplay between different regulatory processes in OA pathogenesis. Potential approaches for these investigations could include CRISPR-Cas9 mediated modulation of splice sites, minigene assays to confirm splicing effects, and functional assays to assess the impact on chondrocyte biology and OA-related processes.

Despite these limitations, our study represents a significant leap forward in understanding the role of alternative splicing in OA. To our knowledge, this is the first sQTL analysis in chondrocytes, coupled with colocalization analysis and functional validation. The identification of novel risk genes and splicing events provides potential new targets for therapeutic intervention and biomarker development in OA.

In conclusion, our study highlights the critical importance of splicing regulation in OA pathogenesis and provides a valuable resource for future investigations. As we continue to unravel the complex genetic and molecular underpinnings of OA, integrative approaches that consider multiple layers of gene regulation will be instrumental in developing more effective diagnostic tools and targeted therapies for this debilitating disease.

## Methods

### Dataset and sample information

The RNA-seq data from primary human chondrocytes were initially collected and processed as described in Kramer et al.^42^.The primary dataset, comprising 202 samples from 101 non-osteoarthritis (non-OA) donors (101 PBS control and 101 FN-f stimulated), is available in the Database of Genotypes and Phenotypes (dbGaP, accession: phs003581.v1.p1, as of June 2024). Our analysis preceded the dbGaP submission. Human talar cartilage tissue was obtained from 101 deceased donors without known arthritis history (Gift of Hope Organ and Tissue Donor Network-https://giftofhope.org/ through the Rush Medical Center, Chicago IL.). We additionally acquired chondrocytes from 16 donors undergoing knee joint replacement due to OA. Chondrocyte isolation and culture methods followed previously established protocols^53^. Cells were treated with phosphate-buffered saline (PBS) as a control or 1 μM recombinant human fibronectin fragment (FN7-10; aka FN-f) for 18 hours each, simulating OA-like conditions as previously described^42^. OA samples underwent identical procedures, excluding FN-f treatment. DNA extraction employed the QIAamp DNA mini kit (Qiagen, #51304), with genotyping performed using the Infinium Global Diversity Array-8 v.10 Kit (Illumina #20031669). RNA extraction utilized the RNeasy kit (Qiagen #74104) with on-column DNase digestion. RNA integrity assessment used the Agilent TapeStation 4150. The New York Genome Center conducted RNA-seq library preparation and sequencing. Genotype processing and quality control for the 101 non-OA samples followed previously described methods^42^.

### RNA-seq processing and quality control

We reprocessed RNA-seq data using updated references, following our previous study’s methodology^42^. Libraries averaged approximately 101 million paired-end reads (2 × 100bp) per sample. Low-quality reads and adapters were trimmed using TrimGalore! (v0.6.7)^54^, followed by FastQC (v0.11.9)^55^ quality control. Trimmed fastqs were aligned to the GENCODE release 45 (GRCh38.p14) reference genome using STAR aligner (v2.7.7a)^56^. Gene expression levels were estimated using Salmon (v1.9)^57^ with --seqBias, --gcBias, and --validateMappings flags, assembly of GENCODE version 45 transcript sequences. The tximeta^58^ R package facilitated gene-level scaled transcript analysis. RNA signal tracks for PBS resting, FN-f stimulated chondrocytes, and OA donors were created using deepTools (v3.5.4)^59^ and merged by condition. We evaluated genotype consistency between RNA-seq and genotyping array data using VerifyBamID (v1.1.3)^60^. Two genotyping sample swaps were corrected. Samples with FREEMIX and CHIPMIX scores > 0.2 post-correction were omitted. This process was applied to OA donor samples as well. The final dataset comprised 101 resting and 101 FN-f stimulated chondrocyte samples from non-OA donors, and 16 OA donor samples.

### Splicing Quantification of intron usages

We quantified alternative splicing events using LeafCutter(v0.2.9)^61^, which allows for the detection of both annotated and novel splicing events. Exon-exon junctions were extracted from uniquely mapped reads using RegTools (v1.0.0)^62^ with default settings. LeafCutter analysis was performed with a minimum threshold of 100 supporting reads per intron and a maximum intron length of 500,000 base pairs, using the GENCODE release 45 (GRCh38.p14) reference genome and GTF as reference. LeafCutter defines intron clusters representing alternative splicing choices by grouping overlapping introns. We calculated the percent spliced in (PSI) for each intron based on its relative usage within its cluster. Intron excision ratios underwent standardization across individuals and quantile normalization across samples. Global alternative splicing principal components were calculated using the prepare_phenotype_table.py script from LeafCutter, utilizing raw counts of intron excision from the intron clustering step.

### Differential splicing analysis

Differential splicing analysis was performed using leaf-cutter_ds.R, considering only introns used in at least 10% of samples and with at least 10 samples per group having sufficient coverage. We compared PBS-control vs. FN-f-stimulated chondrocytes and PBS-control vs. osteoarthritic (OA) chondrocytes. PSI values were normalized using a custom method that accounts for cluster-wise proportions and handles missing data. The batch correction was determined by correlations between technical confounders and the top 10 principal components of global splicing in resting and FN-f stimulated chondrocytes. We identified that the RNA extraction kit batch and fragment batch were the primary sources of batch effects in our data. To address these issues, along with donor-specific variations, we applied the removeBatchEffect() function from the limma package^15^. For the final differential splicing analysis, we employed a mixed-effects model:

model.matrix(∼condition + covariates + (1|DonorID)).

Covariates included RNA shipping date, RNA extraction kit batch, and fragment batch. Significantly spliced intron junctions were identified using a threshold of adjusted p-value (padj) < 0.05 (Benjamini-Hochberg procedure for multiple test correction) and absolute PSI difference (|ΔPSI|) > 0.15.

### Characterization of differentially spliced genes

Significantly spliced intron junctions (padj < 0.05 and |ΔPSI| > 0.15) were identified in comparisons of PBS-control vs. FN-f stimulated chondrocytes and PBS-control vs. OA chondrocytes. We performed pathway (KEGG, Reactome source) and Gene Ontology (GO) analyses on the associated genes using the findMotifs.pl script from the HOMER^63^ software suite (v4.11). Enriched GO terms (p < 0.01) and KEGG and Reactome pathways (p < 0.01) were identified, with GO terms further refined based on semantic similarity using rrvgo (v1.14.2)^64^. Protein-protein interaction networks were constructed using the STRING database (v12.0)^65^, considering only high-confidence interactions (combined score > 0.9).

### Comparative analysis of splicing in OA and FN-f stimulated chondrocytes

We compared differentially spliced intron junctions associated genes between PBS-control vs. FN-f stimulated and PBS-control vs. OA chondrocytes. Both comparisons used the same thresholds (padj < 0.05 and |ΔPSI| > 0.15). To assess the relationship between OA-associated and FN-f-induced splicing changes, we examined the ΔPSI in OA for genes showing differential splicing in response to FN-f in chondrocytes. The significance of these comparisons was evaluated using Wilcoxon tests, with p < 0.01 considered significant.

### sgRNA design and RNP complex formation

Two custom Alt-R crRNAs targeting SNRNP70 (5’-TTGGTT-GAGGACAGCCCCCT-3’ and 5’-TATCACCCCACTGCCCGCTG-3’) were designed to target the plus and minus strands, respectively. These crRNAs, synthesized by Integrated DNA Technologies (IDT), exclude the PAM sequence. Guide RNA (gRNA) duplexes were formed by combining 1.5 μL each of Alt-R tracrRNA (IDT, cat. 1073190) and crRNA in PCR tubes. The mixtures were annealed at 95°C for 5 minutes and cooled to room temperature. Ribonucleoprotein (RNP) complexes were assembled immediately before nucleofection by combining 1.2 μL of annealed gRNA with 2.6 μL of Alt-R Cas9 Nuclease V3 (IDT, cat. 1081058).

### Nucelofection of primary human chondrocytes

Primary human chondrocytes isolated from macroscopically normal cartilage of cadaveric ankle were cultured in DMEM/F-12 medium supplemented with 10% FBS and antibiotics. Approximately 3 × 10^5^ cells per condition underwent nucleofection using the Lonza Nucleofector system (program ER100). Cells were resuspended in 16 μL nucleofector solution, combined with 5 μL RNP complex (or PBS for mock control) and 1 μL Cas9 Electroporation enhancer (IDT, cat. 1075916), then electroporated. Post-nucleofection, cells were cultured in an antibiotic-free medium with 20% FBS at 37°C and 5% CO2 for at least one week.

### Genome editing and colony screening

Genomic DNA was isolated using a column-based extraction method, following a protocol from the QIAGEN DNeasy Blood & Tissue Kit protocol (QIAGEN, Cat. 69504). PCR screening targeted the SNRNP70 alternative exon 8 region using specific primers (Forward: 5’-CTCCTCCCTCTGTTTCT-GATG-3’, Reverse: 5’-CAGGAAAGGGGAGTCGTAGAG-3’). PCR reactions (20 μL total) contained 1-5 μL genomic DNA, 10 μL Platinum II hot-start Green master mix (Invitrogen, cat 14001012), and 0.5 μL each of forward and reverse primers (10 μM stocks). PCR cycling conditions were: 94°C for 2 minutes; 40 cycles of 94°C for 15s, 57°C for 30s, 68°C for 45s; final extension at 68°C for 2 minutes. Products were analyzed by 1.5% agarose gel electrophoresis stained with SYBR Safe DNA Gel stain (Thermo Scientific, cat. S33102).

### Single-cell colony selection and expansion

Edited cells were seeded at 200 cells per 6 cm^2^ dish. After 20 days, individual colonies were manually isolated. Selected colonies were expanded from 24-well plates and plated as passage 1 cells. As previous paper described^66^, cells were cultured in media supplemented with 1 ng/mL TGF-β1(Life Technologies, cat.PHG9214) and 5 ng/mL bFGF(Life Technologies, cat.PHG0264). All 46 selected single colonies were confirmed edits previously described above method section. Further*SNRNP70* alternative exon 8 deletion was confirmed by Sanger sequencing of PCR products from both wild-type and potential heterozygous knockdown (Het-KD) clones. For Het-KD clones, the 512 bp band was gel-extracted. PCR products were purified using Exonuclease I (EXO1) and Shrimp Alkaline Phosphatase (SAP) treatment, following Eurofins Genomics sample preparation guidelines. Purified products were sequenced using SimpleSeq tubes (Eurofins Genomics: https://eurofinsgenomics.com/). Five wild-type clones were combined into one group, and four heterozygous knockout clones into another. Each group was plated into 6 wells: two for sequencing, one as backup, and two for RNA-seq experiments.

### RNA sequencing of Het-KD cells

Cells were expanded to 500,000 cells per well in six-well plates. Total RNA was extracted from two sequencing replicates of each cell population using the RNeasy kit (Qiagen #74104) with on-column DNase digestion. RNA quality and quantity were assessed using the Qubit RNA High sensitivity assay (Thermo Fisher Scientific #Q32582) and Agilent TapeStation 4150 system. Library preparation was performed using the KAPA RNA HyperPrep Kit with RiboErase (Roche #KK8560) following the manufacturer’s protocol. Libraries were quantified using the KAPA Library Quantification Kit (Roche #07960298001). Sequencing was conducted on a NextSeq 2000 system using NextSeq™ 1000/2000 P1 Reagents (100 cycles, Illumina #20074933), generating an average of 24.5 million paired-end reads (2 × 50 bp) per sample.

### Edited cell quantification and differential gene expression analysis

RNA-seq data from edited cells were processed using a pipeline similar to the one described for the PBS-control, FN-f stimulated, and OA chondrocytes RNA-seq data. Briefly, quality control with FastQC (v0.11.9)^55^ and reads were aligned to the GENCODE release 45 (GRCh38.p14) reference genome using STAR aligner (v2.7.7a)^56^. Gene expression levels were quantified using Salmon (v1.9)^57^ with --seqBias, --gcBias, and --validateMappings flags, utilizing GENCODE version 45 transcript sequences. Gene-level scaled transcript analysis was facilitated by the tximeta^58^ R package. Differential gene expression analysis between wild-type and *SNRNP70* alternative exon 8 heterozygous knockout cells was performed using DESeq2^67^. We filtered out lowly expressed transcripts, retaining only those with at least 10 counts in 10% or more of the samples. Differential expression was defined by an FDR-adjusted p-value < 0.01 and absolute log2 fold change > 2, using the apeglm method for fold change shrinkage.

### *SNRNP70* Exon-level quantification

Exon-level counts for *SNRNP70* were generated using featureCounts (subread v2.0.6)^68^. The resulting counts were analyzed using the EBSEA (v1.33.0)^69^ R package, which performs statistical testing at the exon level before aggregating results to the gene level. This approach increases statistical power while accounting for exon dependence using empirical Brown’s method^70^. We considered exons with p-value < 0.05 as significantly differentially used between wild-type and heterozygous knockout samples.

### Characterization of edited cells

Differentially expressed genes (log2FC > 2 and FDR-adjusted p-value < 0.01) were analyzed for KEGG and Reactome pathway enrichment, as well as Gene Ontology (GO) terms. Genes were split based on up- or down-regulation in heterozygous knockout cells compared to wild-type. Enriched pathways and GO terms (p < 0.01) were identified using the findMotifs.pl script from HOMER (v4.11)^63^. GO terms were further refined based on semantic similarity using rrvgo (v1.14.2) with the following parameters:

simThresh=0.7, simMethod=“Rel” ^71^.

### Comparative analysis with FN-f stimulation

We compared the differential genes between *SNRNP70* alternative exon 8 heterozygous knockout cells and controls to those observed in FN-f stimulated chondrocytes (from our previous PBS-control vs. FN-f stimulated analysis). Log_2_ fold changes were examined for genes showing differential expression in both conditions. The significance of these comparisons was evaluated using Wilcoxon tests, with p < 0.01 considered significant.

### Splicing Quantitative Trait Loci (sQTL) Analysis

We performed sQTL analysis to identify genetic variants associated with alternative splicing events in chondrocytes under resting (PBS) and osteoarthritis-like (FN-f stimulated) conditions. This analysis was built upon the differential splicing analysis described earlier, using the same PSI values derived from LeafCutter^61^. We tested genetic variants within ±100 kb of each intron cluster for association with PSI values. This window size was chosen based on previous studies that have demonstrated the enrichment of sQTL SNPs in proximal regions of splice sites^72^. The ±100 kb range allows for the capture of both proximal and distal regulatory elements that may influence splicing^73^. We applied stringent filtering criteria for variant inclusion using PLINK (v1.90b3) to ensure robust associations^74^. We retained only variants with at least two heterozygous donors, no homozygous minor allele donors, or at least two minor allele homozygous donors for autosomal chromosomes. We also required at least two haploid allele counts for the X chromosome. To account for population structure, we calculated genotype principal components (PCs) using bcftools (v1.10) and PLINK (v1.90b3)^74,75^. Our models included the first 10 PCs, explaining approximately 60% of the genetic variance as covariates. We employed QTLtools (v1.3), which extends FastQTL functionality, for sQTL mapping^28^. We incorporated covariates to control for known technical variables and hidden factors, optimizing selection through iterative testing of global splicing principal components (PCs) 1-20. The final sQTL models were:

For PBS samples: *PSI ∼ SNP + 10 PCs of genotype + 5 PCs of global splicing + FragmentBatch + RNAextractionKitBatch + SequencingBatch*

For FN-f samples: *PSI ∼ SNP + 10 PCs of genotype + 4 PCs of global splicing + FragmentBatch + RNAextractionKitBatch + SequencingBatch*

To determine significant associations, we implemented a permutation-based approach using QTLtools^28^. We performed 1,000 permutations for each phenotype to characterize the null distribution of associations empirically. This permutation scheme approximates the tail of the null distribution using a beta distribution. We used the qtltools_runF-DR_cis.R script to implement the Storey & Tibshirani False Discovery Rate (FDR) procedure, identifying significant sQTLs at 5% FDR^76^. The resulting nominal p-value threshold range was 9.99e-6 to 0.0001 for both PBS and FN-f conditions. To identify multiple independent sQTLs per splice intron junction, we performed conditional analysis using QTLtools^28^. This method uses a stepwise linear regression approach to discover additional independent signals while accounting for multiple testing. The process involves establishing a nominal p-value threshold for each phenotype, iteratively identifying the most robust associations, and assigning nearby variants to independent signals.

### Condition-specific and Response sQTL Analysis

To assess condition-specific effects and potential genetic interactions, we applied inverse normal transformation of PSI ratios. This transformation was necessary to normalize the distribution of PSI values and reduce the impact of outliers, ensuring robust statistical analysis. We then conducted interaction testing using linear mixed-effects models, comparing reduced models (without interaction terms) to full models (including interaction terms) for each sQTL-splice intron junction pair. To investigate genetic variants influencing splicing in a condition-specific manner, we constructed two linear mixed models using the lme4 package^29^ in R:

Null hypothesis: *PSI ∼ genotype + covariates + condition + (1*|*DonorID)*

Alternative hypothesis: *PSI ∼ genotype + covariates + condition + genotype:condition + (1*|*DonorID)*

*H*0: *expression ∼ SNP* + *covariates* + *condition* + (1|*Donor*) *H*1: *expression ∼ SNP* + *covariates* + *SNP:condition* + (1|*Donor*)

Where covariates included the same technical factors and PCs used in the standard sQTL mapping, the condition was coded as 0 (PBS) or 1 (FN-f), and (1|DonorID) accounted for donor-specific random effects. We assessed the significance of the *genotype:condition* term using ANOVA, with p < 0.05 considered significant for the interaction effect.

To identify high-confidence condition-specific QTLs, we applied additional filters: 1)sQTLs detected only in one condition (PBS or FN-f), 2) At least 5 donors per minor allele to ensure robust estimation, 3) Absolute beta difference ≥ 0.2 between conditions, indicating a substantial change in effect size.

### RNA binding protein motif analysis

We selected 140 RNA-binding proteins (RBPs) from the ENCODE project, focusing on all eCLIP-seq data generated for K562, HepG2, and SM-9MVZL cell lines^77^. The eCLIP-seq data were processed, aligned to the GRCh38 genome assembly, and provided as bed format. We used the Functional enrichment of molecular QTLs (Fenrich) module from QTLtools to evaluate the overlap between sQTLs and RBP binding sites^28,78^. A permutation scheme was employed to maintain the distribution of functional annotations and molecular QTLs around molecular phenotypes. We performed 1,000 permutations to estimate the probability of the observed overlaps of the lead sQTLs relative to expectation. For sQTLs overlapping with RBP binding sites, we calculated the Pearson correlation between the RBP gene expression (from the same donor RNA-seq data) and the spliced intron junction usage pattern across all samples. We considered correlations with R^2^ > 1.5 and p-value < 0.05 as significant. we performed KEGG and Reactome pathway enrichment analysis on the genes associated with sQTLs that overlap RBP binding sites and show a significant correlation with RBP expression. using the findMotifs.pl script from the HOMER software suite (v4.11), the KEGG and Reactome pathways (p < 0.01) were identified.

### Colocalization between sQTL and OA GWAS

We conducted colocalization analysis between our sQTL results and OA GWAS signals. Summary statistics for 11 OA phenotypes were obtained from Boer et al., encompassing 826,690 individuals of European and East Asian descent (Musculoskeletal Knowledge Portal, https://msk.hugeamp.org/). These statistics were converted to hg38 coordinates for consistency with our analysis. Overlapping signals were identified by selecting lead sQTL variants in moderate linkage disequilibrium (LD, r^2^ > 0.5) with lead GWAS variants. LD was calculated using PLINK (v1.90b3.45) with parameters --ld-window 200000 --ld-window-kb 1000, based on the 1000 Genomes European reference panel phase 3, chosen due to the East Asian population sample size > 4,000 individual^4,74,79^. We defined a window encompassing the sQTL and GWAS lead variants for each pair of overlapping signals, extending by 250 kb in each direction. To ensure consistent identification, we mapped variants to rsIDs using dbSNP155, querying dbSNP VCF files with bcftools, and custom scripts to match variants based on chromosome, position, and alleles. Colocalization analysis was performed using the Bayesian method *coloc* package, implemented in R^80^. The coloc.abf function was used to calculate posterior probabilities for various hypotheses, including H3 (association with sQTL only) and H4 (association with both sQTL and GWAS traits). Signals with a posterior probability for H4 (PP H4) > 0.7 were considered as evidence of significant colocalization. We also calculated the ratio (PP H4)/(PP H3 + PP H4) to quantify the strength of colocalization evidence.

This analysis was conducted separately for sQTLs identified in PBS and FN-f conditions to examine condition-specific effects on OA risk.

### Visualization

Protein-protein interaction networks were visualized using Cytoscape(v3.9)^81^. Heatmaps were generated using the ComplexHeatmap R package(v3.19)^82^. Genomic signal tracks and locus zoom plots were created with plotgardener (v1.9.4). All other plots were produced using ggplot2 in R^83^. These visualizations were generated using R version 4.3.

## Data Availability

The genotyping and RNA-seq datasets generated for this study, including 101 non-OA samples, 16 OA samples, and edited cell RNA-seq data, are available in the NIH’s database of Genotypes and Phenotypes (dbGaP) under accession number phs003581.v1.p1 (data submission in process).

## Code Availability

All analysis code used in this study is available at https://github.com/seyoun209/CQTL_sQTL

### Acknowledgments

This work was supported by NIH grants (R01AR079538 to D.H.P. and R.F.L., R35-GM128645 to D.H.P., R37-AR049003 to R.F.L., R21-AR084104 to B.O.D.) and training grants (1F31AR083722-01A1 to S.B., and T32GM135123-02 to S.B.). The project was also supported by the National Center for Advancing Translational Sciences (NCATS) through NIH Grant UL1TR002489 and by the UNC Thurston Arthritis Research Center through a pilot and feasibility grant. This study was also supported by Rush University Klaus Kuettner, Chair for Osteoarthritis Research (S.C.).

We would like to thank the University of North Carolina at Chapel Hill and the Research Computing group for providing computational resources and support that have contributed to these research results. We thank Jason Stein, Nil Aygün and Jordan Valone for their guidance and assistance with sQTL analysis and Erika Deoudes for her graphic design contributions. We also thank the Gift of Hope Donor Network and the donor families for providing the normal donor tissue and the UNC Department of Orthopedics for the OA tissue.

## Author information

### Contributions

S.B., R.F.L., B.O.D., and D.H.P. conceived the project and designed the study. P.C. conducted the FN-f treatment and collected all the cells. S.C. assisted with all the donor collection. S.D. and E.T. performed experiments on all of the RNA-seq and genotyping samples. N.K. processed quality control of all RNA-seq and genotyping data. S.B. processed and analyzed the genomic data and was supported by R.F.L., B.O.D., and D.H.P. S.B. and J.S. performed all edited cell experiments. S.B., R.F.L., B.O.D., and D.H.P. prepared the first draft of the manuscript. All authors contributed to the final manuscript.

### Corresponding authors

Correspondence to Richard F Loeser, Brian Diekman, or Douglas H Phanstiel

## Supplemental Information

**Supplemental Fig 1.**
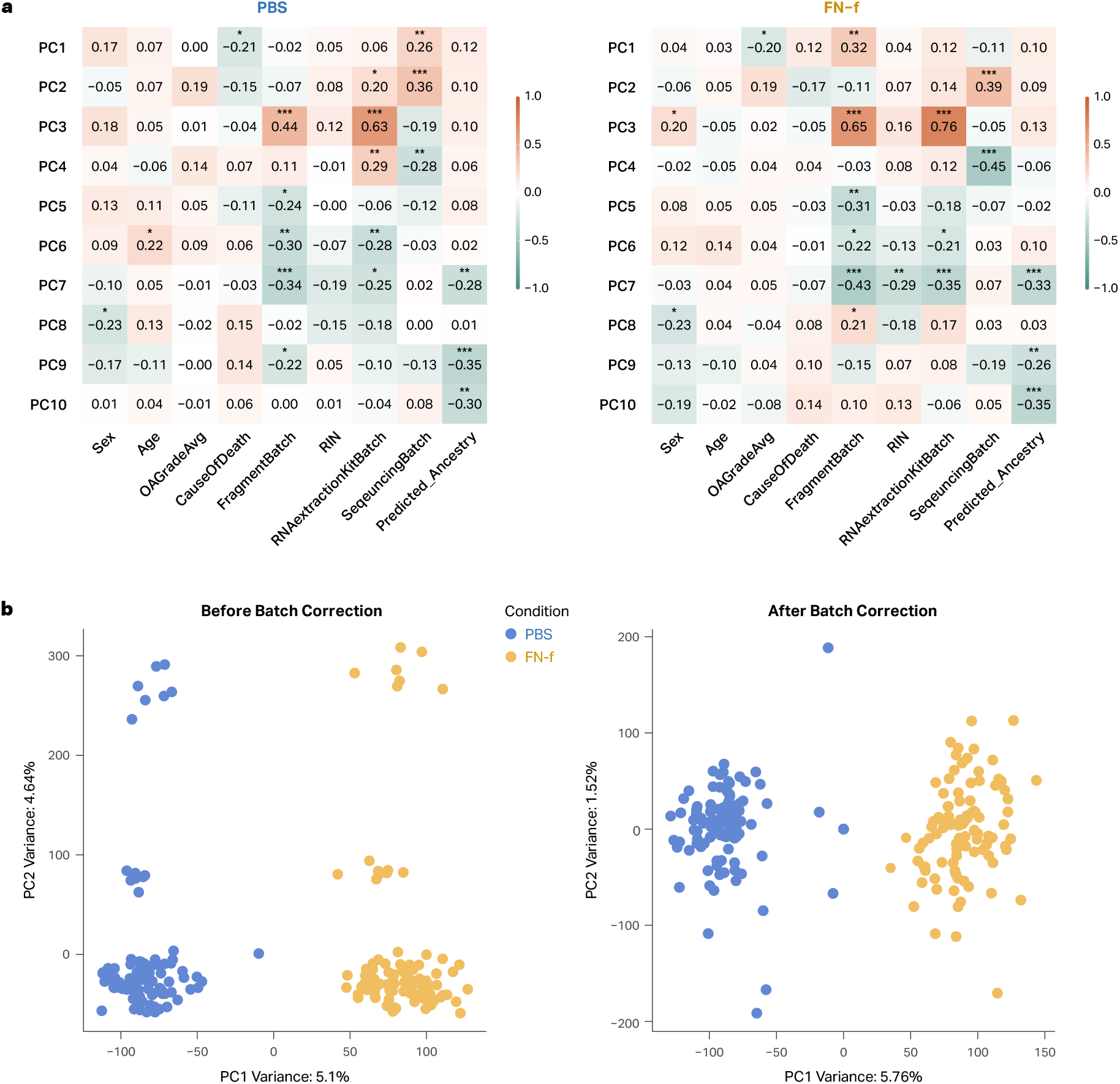
Batch correction of splice ratio. (**a**) Heatmaps showing correlations between technical confounders and top 10 principal components of global splicing in resting (PBS) and activated (FN-f stimulated) chondrocytes (asterisks indicate significant correlations: * p < 0.05, ** p < 0.01, *** p < 0.001). Confounders include: Sex: Biological sex of the tissue donor Age: Age of the deceased tissue donor OAGradeAvg: Average osteoarthritis grade of the cartilage tissue, only normal-appearing tissue (grades 0–1) CauseOfDeath: Various reasons for deceased tissue donors FragmentBatch: Recombinant human FN-f (FN7-10) batch RIN: RNA Integrity Number, a measure of RNA quality RNAextractionKitBatch: Batch effects due to the RNeasy kit performed at different times SequencingBatch: Potential batch effects due to sequencing performed in different runs Predicted_Ancestry: Genetic ancestry based on the 1000 Genomes Project super-populations, determined using EIGENSTRAT on genetic principal components (**b**) Principal Component Analysis (PCA) plots of all spliced intron junctions (n=86,548) before and after batch correction.

**Supplemental Fig 2.**
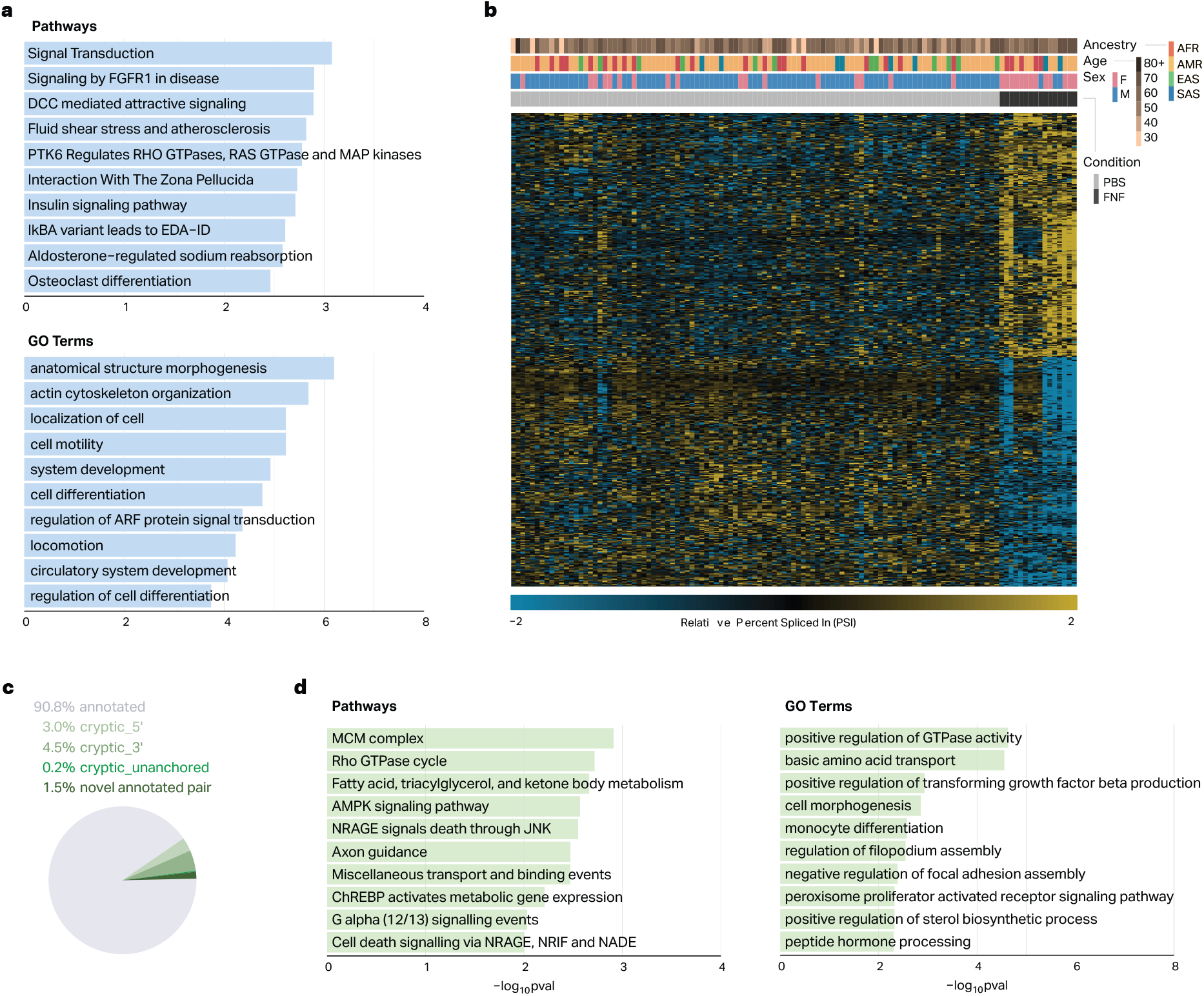
Differential alternative splicing changes in FN-f and OA conditions. (**a**) Barplot of select KEGG & Reactome pathways and GO terms enriched in the 590 differentially spliced genes from the FN-f vs. PBS comparison. (**b**) Heatmap of differential splicing events between PBS control and OA samples. Displaying PSI values for top intron junctions (n = 329) from LeafCutter clusters, representing 321 genes (padj < 0.05, | PSI| > 0.15). (**c**) The pie chart illustrates the distribution of differential alternative splicing intron junction types (n = 466): annotated (n = 423, 90.8%), cryptic five prime (n = 14, 3.0%), cryptic three prime (n = 21, 4.50%), cryptic unanchored (n = 1, 0.2%), and novel annotated pair (n = 7, 1.50%). (**d**) Barplot of select GO terms and KEGG & Reactome pathways enriched in the 321 differentially spliced genes from OA vs. PBS control.

**Supplemental Fig 3.**
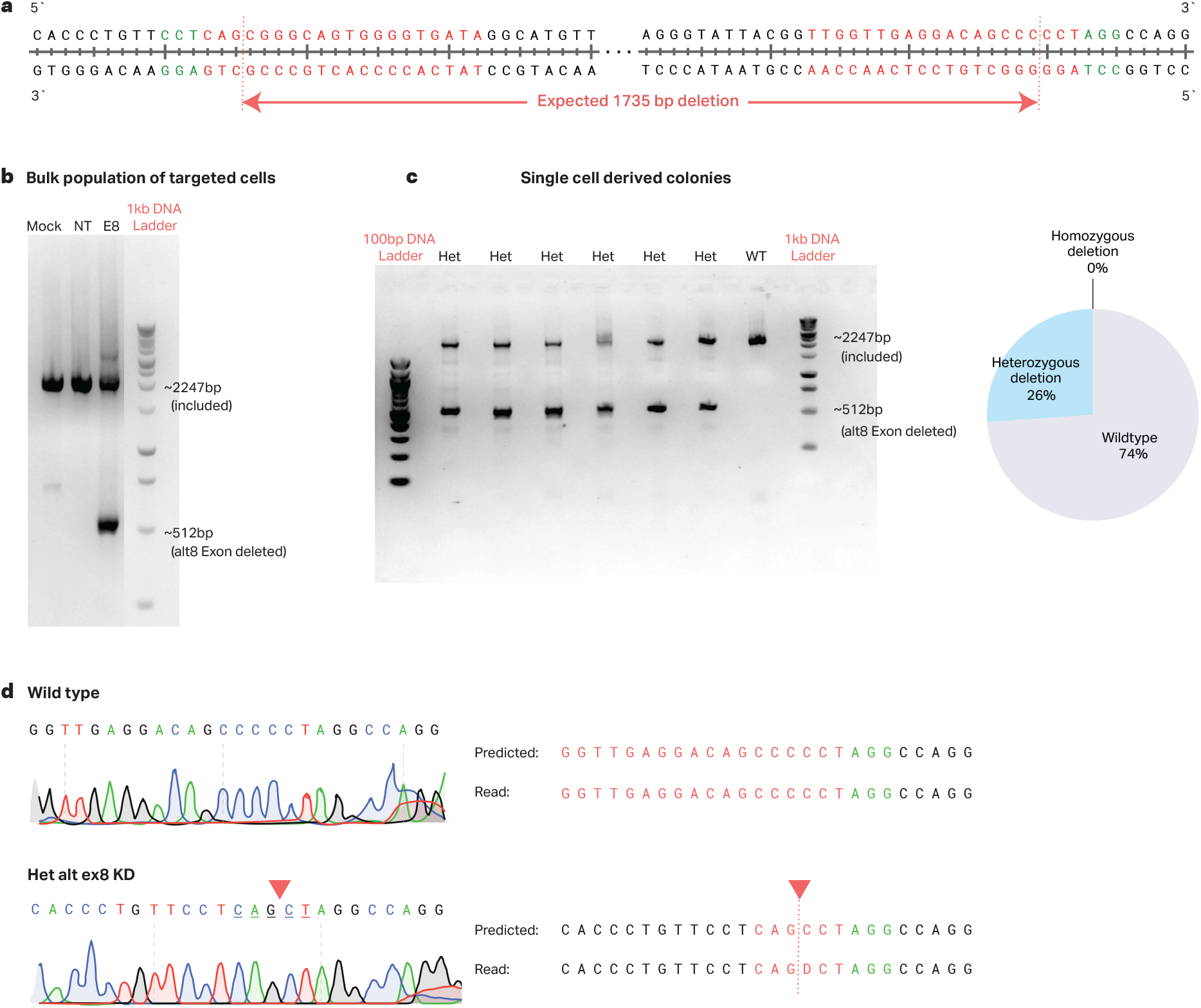
CRISPR/Cas9-mediated targeting of *SNRNP70* alternative exon 8. (**a**) Region of the *SNRNP70* gene sequence containing alternative exon 8 targeted by CRISPR/Cas9 editing. PAM sites are highlighted in green, guide RNA target sites are shown in red, and expected cut sites are indicated by arrows. (**b**) Representative gel electrophoresis of the bulk cell population after targeting. The ∼1735 base pair (bp) shorter product indicates successful *SNRNP70* alternative exon 8 targeting. Lanes represent Mock (no guide RNA), NT (non-targeting control crRNA), and E8 (guide RNAs targeting alternative exon 8). (**c**) Example of single cell-derived colonies showing different targeting outcomes: WT (wildtype, Full-length product) and Het (heterozygous, both Full-length and edited shorter product). No homozygous knockout was observed. Right: Pie chart illustrating the proportion of wildtype (WT) and heterozygous deletion (HET) outcomes in analyzed colonies. Homozygous deletion was not observed (0%). (**d**) Sanger sequencing results from representative colonies. The predicted and actual read sequences for wildtype and heterozygous alternative exon 8 knockdown (Het alt ex8 KD) samples are shown. Red triangles indicate expected cut sites. In the Het alt ex8 KD sample, the actual sequence shows a deletion of one cytosine (C) beyond the expected cut site. The dash (-) in the Het alt ex8 KD read sequence represents the single base pair deletion.

**Supplemental Fig 4.**
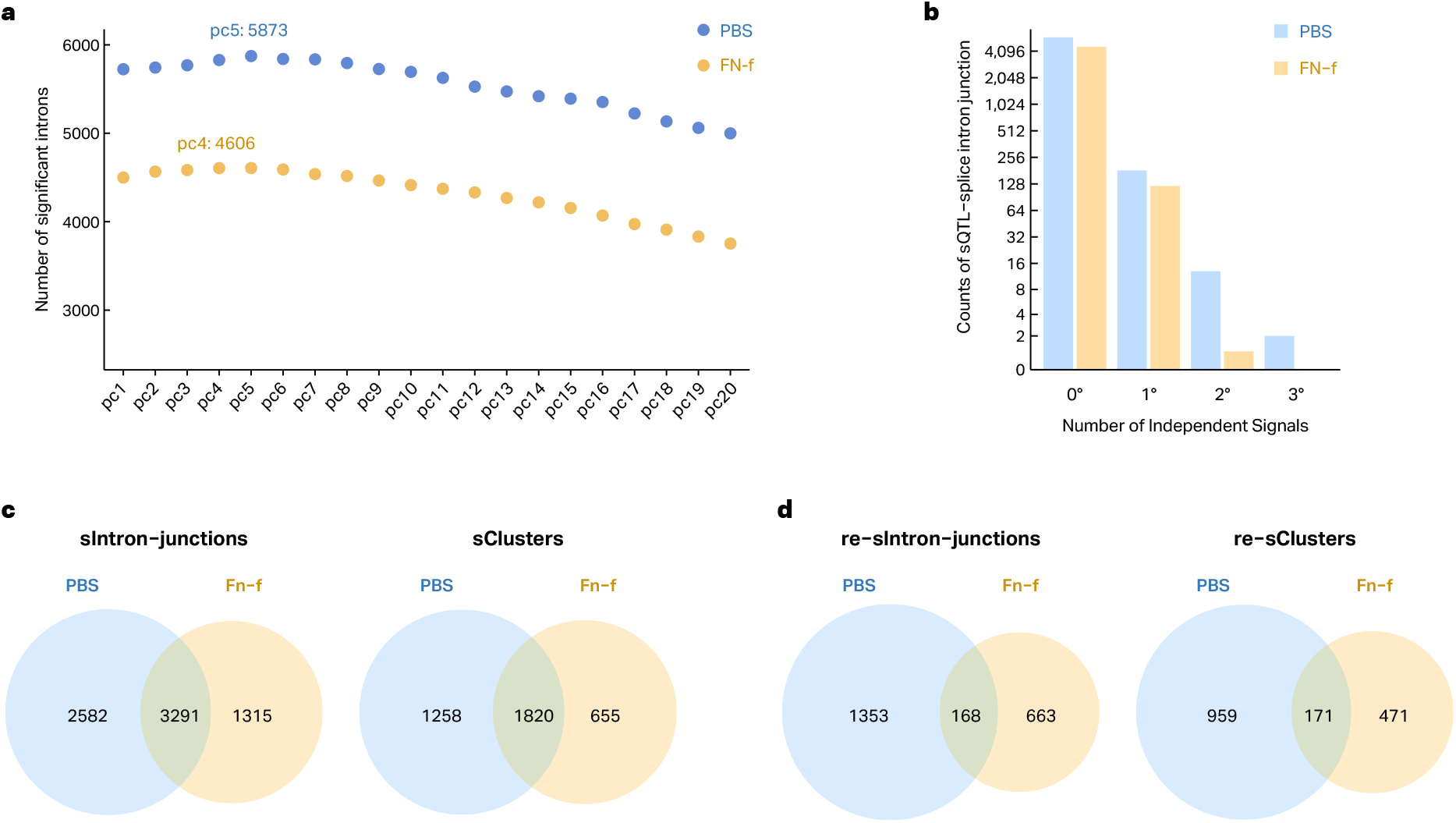
Genetic differences impact differential splice intron junction in resting and activated chondrocytes. (**a**) Covariate selection analysis for sQTL using QTLtools. Dots represent significant splice introns for global splicing PCs 1-20. Blue: PBS (resting), yellow: FN-f (stimulated). Y-axis: number of significant splice intron junctions after permutations. PBS PC5 (n=5873) and FN-f PC4 (n=4606) identified the most sQTL-splice intron junction pairs. (**b**) Barplot showing the number of sQTL-splice intron junctions (y-axis, log2 scale) identified by conditional analysis (x-axis). Primary signal (0°) and additional signals (1° to 3°) are shown. PBS control is blue, and FN-f stimulated is yellow. Only the PBS condition has quaternary (3°) signals. (**c**) Left: Venn diagram illustrating the overlap of sQTL-splice intron junction pairs identified in PBS and FN-f stimulated conditions. Right: Venn diagram showing the overlap of sQTL pairs associated with clustered splice introns. Clustered splice introns represent groups of splice junctions sharing the same Group ID. (**d**) Left: Venn diagram depicting condition-specific and genetic interaction (response) sQTL-splice intron junction pairs identified in PBS and FN-f stimulated conditions. Right: Venn diagram showing the overlap of response sQTLs associated with clustered splice introns. As in (c), clustered splice introns are defined by shared Group IDs.

**Supplemental Fig 5.**
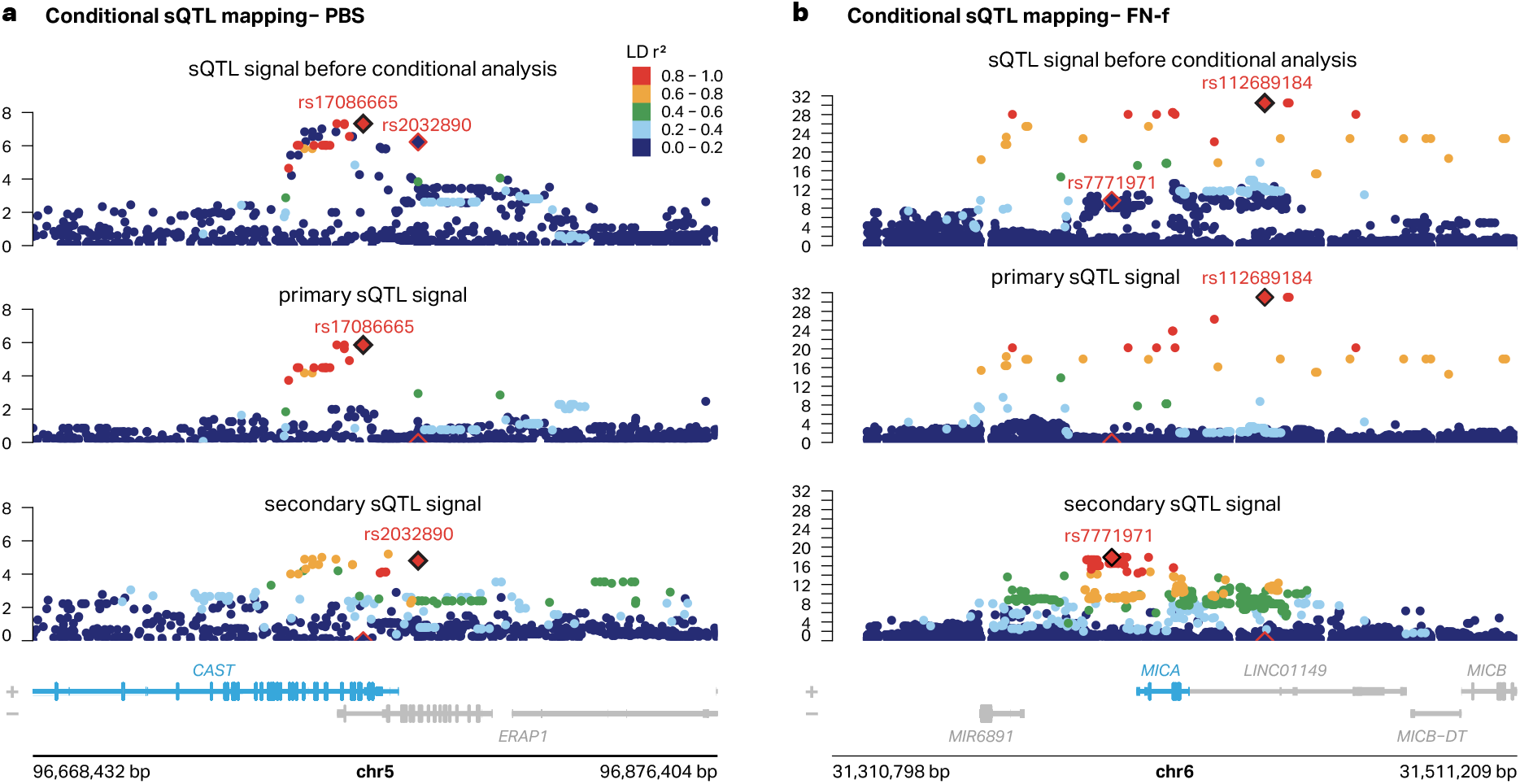
Conditional sQTL mapping identifies secondary signals. (**a**) Example of the PBS (resting) condition for *CAST* splice intron junction (chr5:96768431-96776404). Top: Locus zoom plot before the conditional analysis. Middle: Locus zoom plot of the primary signal (rs17086665). Bottom: Locus zoom plot of the secondary signal (rs2032890). Each signal is colored by LD r^2^ relative to the signal lead variant. (**b**) Example of the FN-f (stimulated) condition for *MICA* splice intron junction (chr6:31410797-31411209). Top: Locus zoom plot before conditional analysis. Middle: Locus zoom plot of the primary signal (rs112689184). Bottom: Locus zoom plot of the secondary signal (rs7771971). Both variants were not in LD with each other.

